# Accurate 4Pi single-molecule localization using an experimental PSF model

**DOI:** 10.1101/2020.03.18.997163

**Authors:** Yiming Li, Elena Buglakova, Yongdeng Zhang, Jervis Vermal Thevathasan, Joerg Bewersdorf, Jonas Ries

**Affiliations:** Department of Biomedical Engineering, Southern University of Science and Technology, Shenzhen 518055, China; European Molecular Biology Laboratory, Cell Biology and Biophysics, Heidelberg 69117, Germany; Moscow Institute of Physics and Technology, Dolgoprudny 141701, Russia; Department of Cell Biology, Yale School of Medicine, New Haven, CT 06511, USA; Faculty of Biosciences, Heidelberg University, Heidelberg 69120, Germany; Kavli Institute for Neuroscience, Yale School of Medicine, New Haven, CT 06511, USA; Nanobiology Institute, Yale University, West Haven, CT 06516, USA; Department of Biomedical Engineering, Yale University, New Haven, CT 06511, USA

## Abstract

Interferometric single-molecule localization microscopy (iPALM, 4Pi-SMS) uses multiphase interferometry to localize single fluorophores and achieves nanometer isotropic resolution in 3D. The current data analysis workflow, however, fails to reach the theoretical resolution limit due to the suboptimal localization algorithm. Here, we develop a method to fit an experimentally derived point spread function (PSF) model to the interference 4Pi-PSF. As the interference phase is not fixed with respect to the shape of the PSF, we decoupled the phase term in the model from the 3D position of the PSF. The fitter can reliably infer the interference period even without introducing astigmatism, reducing the complexity of the microscope. Using a spline-interpolated experimental PSF model and by fitting all phase images globally, we show on simulated data that we can achieve the theoretical limit of 3D resolution, the Cramér-Rao lower bound (CRLB), also for the 4Pi microscope.

---

Single-molecule localization microscopy (SMLM, also known as PALM^1^ or STORM^2^), has pushed optical resolution in microscopy towards the nanometer scale and leads to structural insights into cell biological questions^3–5^. 4Pi-SMLM (also called iPALM^6^, 4Pi-SMS^7^ or 4Pi-SMSN^8^) uses interference of the fluorescence signal detected with two opposing objective lenses to drastically increase the z-resolution. Compared to the conventional single objective 3D SMLM system^9–13^, which only collects a hemispherical wavefront emitted from the sample, the dual-objective system collects the (nearly) complete spherical wavefront, thus increasing the aperture angle and doubling the number of detected photons. Overlaying both detection paths on a beam splitter results in self-interference of individual photons, provided the optical path length difference (OPD) is within the coherence length. Three^6^ or four^7,8^ interference phase images are then simultaneously recorded on separate cameras or different regions on one camera chip and provide a very sensitive readout of a fluorophore’s axial position (**Fig. 1a**). The resulting axial resolution can surpass the lateral resolution and is about 6 - 10 times better than the axial resolution achievable with the single objective system. In theory, 3D sub-10 nm resolution can be achieved with only 250 photons collected in each objective for an individual molecule^14^. In practice, however, this theoretical resolution limit has not been achieved in 4Pi SMLM, mainly due to suboptimal analysis approaches. This is in contrast to single-objective 3D SMLM, where the resolution limit can be practically reached by maximum likelihood estimation (MLE) using a bead-calibrated experimental PSF model^15^.

**Figure 1.**
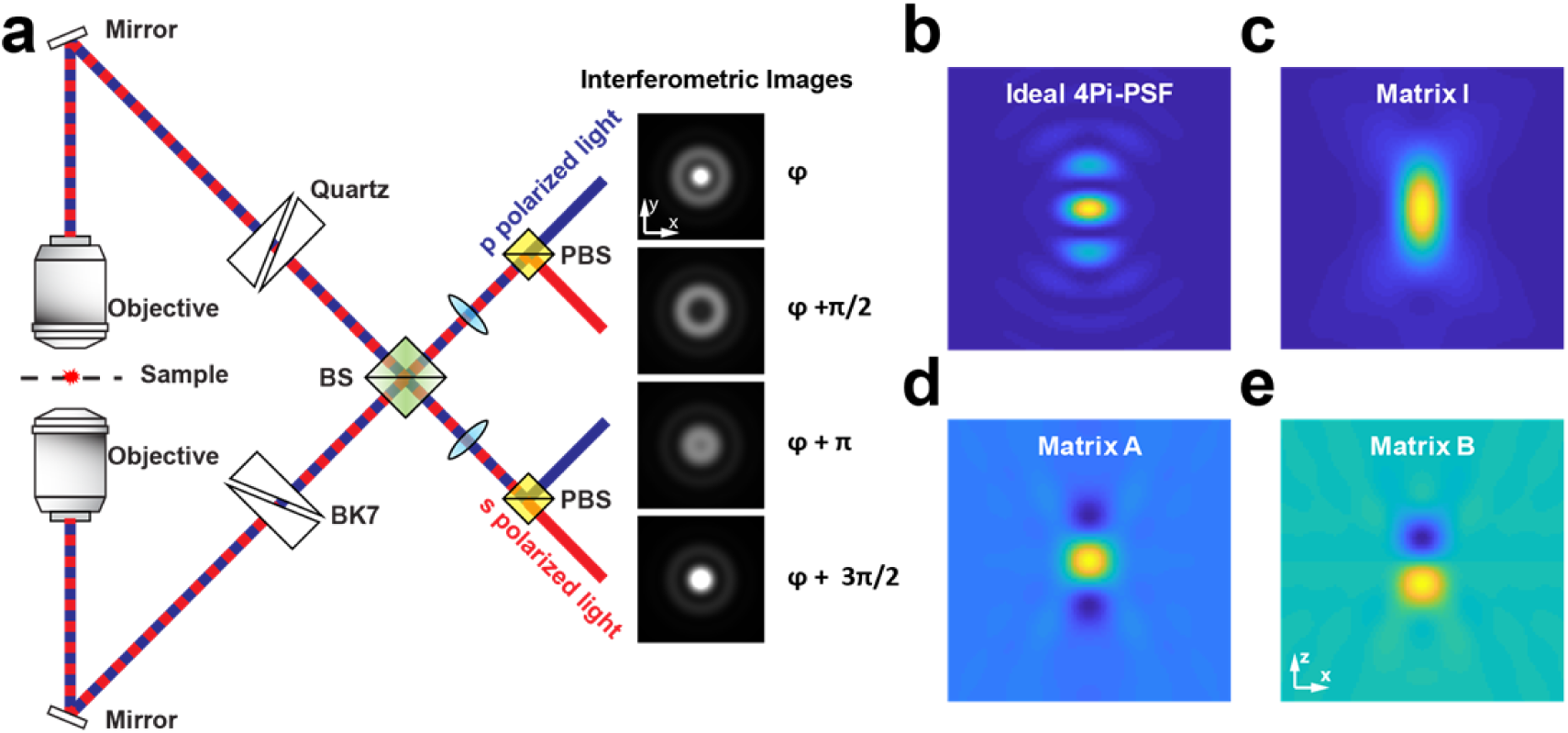
IAB-based 4Pi-PSF model. **(a)** Simplified schematic of a four-phase interference 4Pi-SMLM system. The emission fluorescence is collected by both objectives and interferes with itself at the beam splitter (BS). The phase shift between the p- and s-polarized channel is controlled by two modified Babinet-Soleil compensators (a phase shift of pi/2 is shown). The p- and s-polarized light is separated by polarizing beam splitters (PBS) to create 4 interference images simultaneously. **(b)** Cross section of ideal fully coherent 4Pi-PSF with interference phase of 0. **(c)** Cross section of matrix I, **(d)** of A, **(e)** of B. A full vectorial PSF model^19^ was used for simulations with parameters: NA 1.35. Refractive index 1.40 (immersion medium and sample) and 1.518 (cover glass). Emission wavelength 668 nm. The same parameters are used throughout this work unless noted otherwise.

One of the most commonly used methods to extract the *z*-position from the interference 4Pi-PSF is to determine the phase based on photometry between different interference channels^6–8^. However, this approach cannot extract information from the fringe pattern, thus the localization accuracy is worse than the theoretical limit. Recently, spline interpolated phase retrieved 4Pi PSF models were introduced for fitting of 4Pi-SMLM data^16^. While the method could potentially achieve the theoretical resolution, its usability in practice is limited. The problem is that the interference phase is fixed with respect to the 3D position of the PSF in the phase retrieved 4Pi-PSF model. However, the interference phase can easily drift as even minor changes in temperature will change the OPD in the interference cavity in real experiments. Thus, a simple multi-channel 3D PSF is not well suited to describe real 4Pi-SMLM data.

To overcome these limitations, here we developed a new experimental PSF model that fully describes the 4D nature (*x, y, z* and phase) of the 4Pi-PSF. The OPD between the two interference arms does not only depend on the axial position of the single molecule, but is also affected by the path length of the interference cavity which can change during the experiment. Therefore, we decouple the interference phase from the *z*-position of the point emitter in our new 4Pi-PSF model (IAB model, **Fig. 1b-d, Online methods**). We then interpolate the phase-independent part of the experimental 4Pi-PSF model using cubic splines and globally fit the interference images from all phase channels. For fitting of the parameters globally across different channels, the spatial transformation between different channels can be calibrated using beads on cover glass. Based on whether a parameter is independent in each channel or the same in all channels, the Hessian and Jacobian matrices are constructed by using the information in one channel or across multiple channels^17^. Finally, by iteratively updating the parameters until they are converged, we obtain the maximum likelihood for the Poisson process. We implemented the algorithm on the GPU and reached a fitting speed more than 5 times faster than the CPU fitter (**Supplementary Fig. S1**).

We then investigated the accuracy of our fitting approach on simulated data and compared it to the photometry-based method, as well as the CRLB, which is the limiting lower bound of the variance of estimated parameters for any unbiased estimator. The CRLB was evaluated as the diagonal element of the inverse of the Fisher information matrix. We simulated images of single molecules in 4 phase channels with phase shifts of {0, *π/*2, π, 3*π/*2}, 1000 photons per molecule and a background of 20 photons/pixel for each objective. We simulated 1000 molecules at each axial position (every 50 nm from −600 nm to 600 nm around the focal plane) assuming an astigmatism of 100 mλ. We then evaluated the localization accuracy (root mean square error) of *x, y* and *z*_*φ*_ at different axial positions (**Fig. 2**).

**Figure 2.**
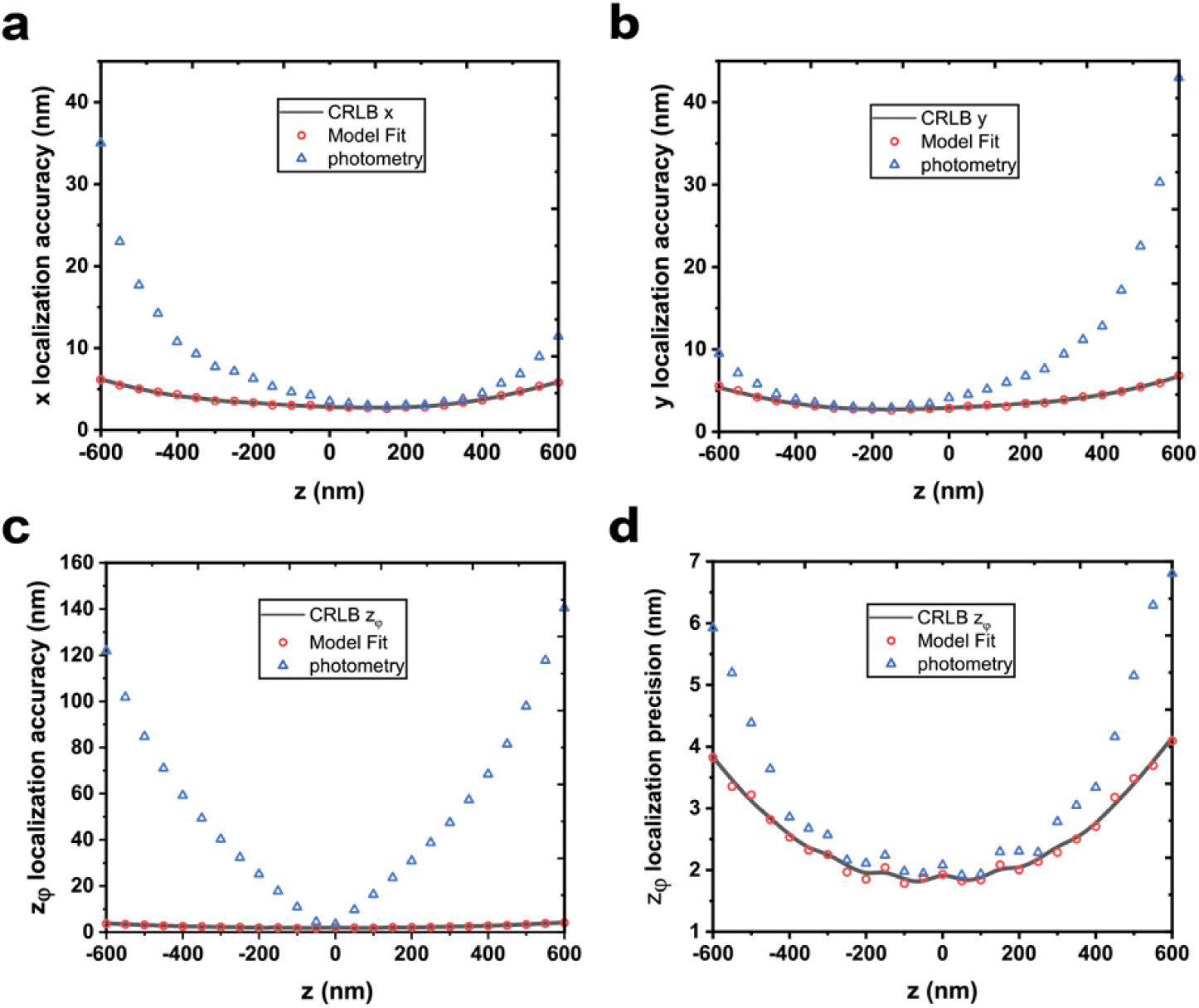
Comparison of localization accuracies. using the IAB-based 4Pi-PSF model fit and the photometry-based analysis method. Localization accuracy in x (a), y (b) and *z*_*φ*_ (c). Localization precision in *z*_*φ*_ (d). The localization accuracy is calculated as the root mean square error and localization precision as the standard deviation.

For the photometry-based method we followed the work by Aquino and coworkers^7^. Here, the interference phase is calculated from the 0^th^ moment of the intensity in the different interference phase channels. The reduced moments are calibrated at each *z*-position and modeled by two sine curves with a known phase shift between p- and s-polarized channels. For each individual single molecule, the interference phase can be analytically solved using the two sine models. The *x* and *y* positions are obtained by fitting the summed image of the all phase channels with a Gaussian model.

Our analysis (**Fig. 2 (a)-(c)**) reveals that the localization accuracy using MLE with our IAB-based 4Pi-PSF model can reach the CRLB in all dimensions. The localization accuracy of dim single molecules (equivalent to 1000 photons and 20 background photons per pixel in single objective system) can be well below 10 nm over a large axial range. In contrast, the localization accuracy for the photometry-based method is worse than the CRLB, specially for molecules away from the focal plane. In *x* and *y*, the improvement from using the new model is up to 4-fold (**Fig. 2 (a)** and **(b)**). The reason is that model fit extracts information from the fringes in the interference PSF, whereas the summed PSF which is used for lateral localization in photometry method does not contain these high spatial frequencies. The large values for the *z* localization accuracy in photometry (**Fig. 2 (c)**) are due to systematic errors, indicating that the intensity of the 0^th^ moment does not follow the sine model. The localization precision (standard deviation) is up to ∼1.5 fold worse (**Fig. 2 (d)**).

As even an unmodified (non-astigmatic) 4Pi-PSF has a different shape in different interference periods^7^, we hypothesized that we can distinguish these interference periods directly using our 4Pi-PSF model fit. This would render the introduction of astigmatism obsolete, greatly simplifying the design of the microscope. To test our hypothesis, we simulated upper and lower objectives PSFs without introducing any aberration and coherently added them to produce an ideal 4Pi-PSF. In order to overcome the problem that the fitter might not converge across interference periods, we explored the bidirectional fitting approach as described before^15^. Here, we fit each single molecule twice with a starting *z*-parameter above the focal plane (300 nm) and another *z*-parameter below the focal plane (- 300 nm). Furthermore, we chose starting phases for these two positions with a phase shift of π. We then chose the solution of these two fits with the maximum likelihood. Similar to the 4Pi-PSF with astigmatism, we also could achieve the CRLB in all directions also for the unaberrated 4Pi-PSF (**Fig. 3**).

**Fig. 3.**
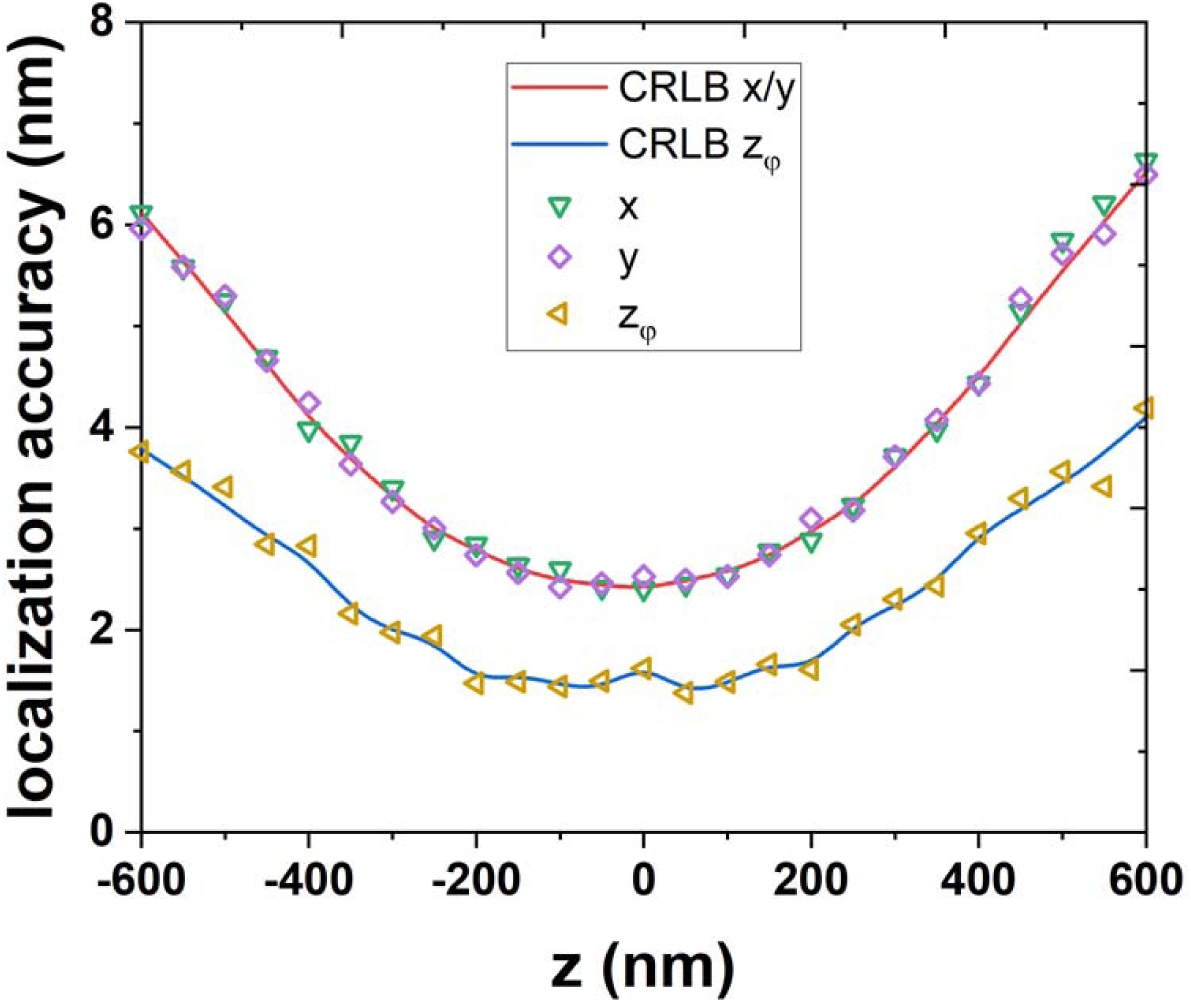
Unaberrated 4Pi-PSF: *x, y* and *z*_*φ*_ localization accuracy of an ideal 4Pi-PSF without additional astigmatism. 1000 4Pi single molecule images were simulated at each *z*-position with 4 phase channels (0, pi/2, pi, 3pi/2). For each objective, 1000 photons/localization and 20 background photons/pixel were used.

To demonstrate that our approach works on real experimental 4Pi-SMLM data, we imaged nuclear pore complexes using a 4Pi microscope with a design based on Huang and coworkers^8^. Nuclear pore complexes are well suited as reference standards as they position the proteins at defined 3D locations in the cell^18^. We used genome edited cell lines in which we labeled Nup96-SNAP with BG-AF647 and embedded the sample in a refractive index matched buffer (**Online Methods**). The two rings of Nup96 are clearly resolved in the axial dimension (**Fig. 4**).

**Fig. 4.**
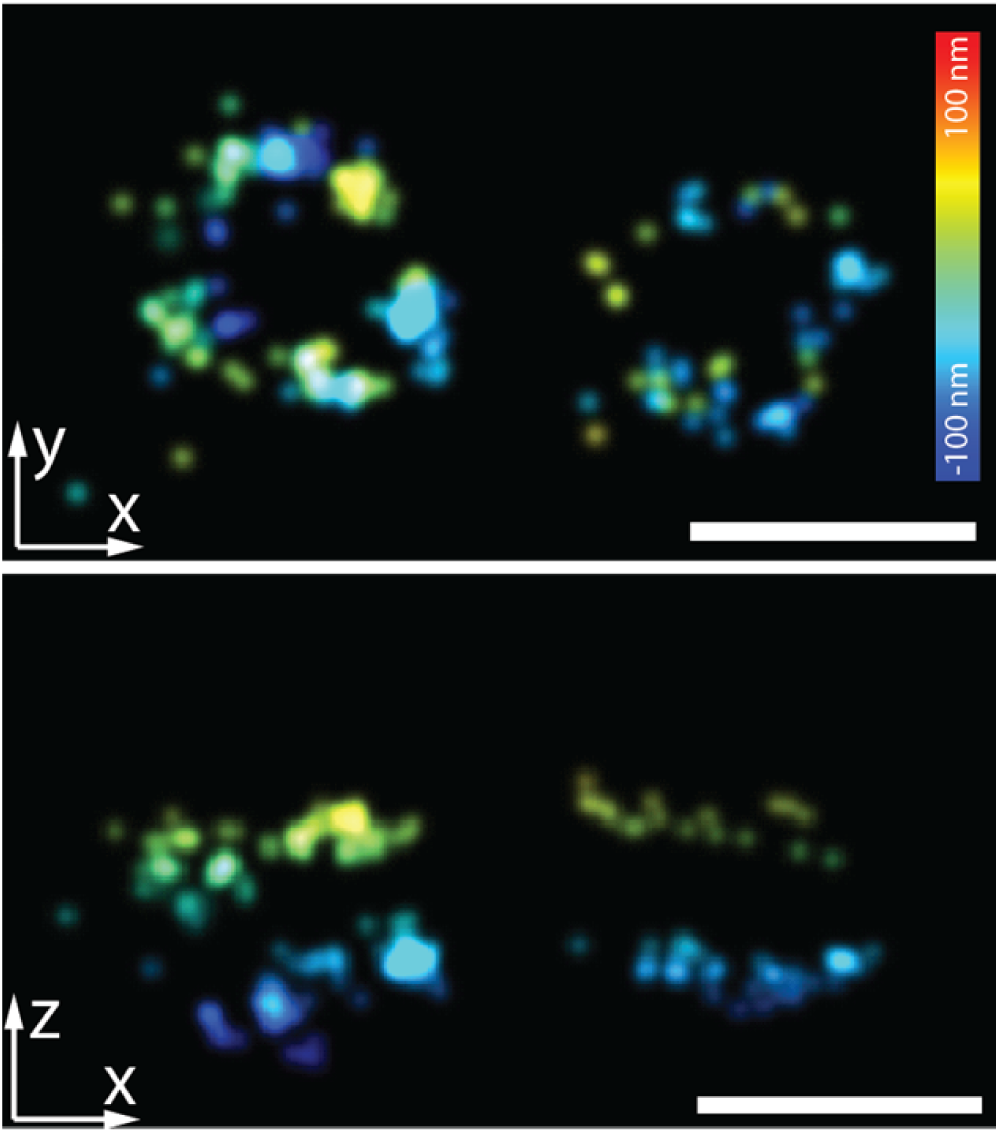
Experimental data on nuclear pore complex. Nup96-SNAP-Alexa647 was imaged within a refractive index matched imaging buffer and reconstructed by the IAB-based 4Pi-PSF model. Scale bars 100 nm.

In conclusion, we developed a new 4Pi-PSF model to fully describe the 4D nature of the 4Pi-PSF. We decoupled the interference phase from the 3D position of the PSF and represent the 4Pi-PSF with three 3D matrices (I, A and B) in combination with a simple phase term. Therefore, instead of calibrating a complete 4-dimensional 4Pi-PSF, we could obtain an experimental 4Pi-PSF just by calibrating the three 3D matrices, which is practically much easier. Furthermore, we developed a complete analysis pipeline to analyze the multi-phase channel 4Pi-PSFs simultaneously. Our global fitter achieved the theoretical minimum uncertainty, the CRLB, in all dimensions. Compared to the conventional photometry-based approach, it greatly improved the localization precision and accuracy, enabling sub-10 nm resolution in all three dimensions even for rather dim single molecule conditions (1000 photons/molecule and 20 background/pixel). Moreover, the IAB-based 4Pi-PSF model fit allowed us to unwrap the phase without introducing additional aberrations that are usually used to distinguish different interference periods, which could simplify the design of 4Pi-SMLM microscopes. Finally, we validated our 4Pi-PSF model by imaging the biological samples, nuclear pore complex protein Nup96, and clearly resolved the double-ring structure with a distance of only 50 nm. We believe that this work is an important step in reaching the full potential of 4Pi-SMLM.

## ACKNOWLEDGEMENTS

We thank P. Hoess for helping with preparation of the refractive index matched buffer. This work was supported by the European Research Council (ERC CoG-724489, J.R.), the EMBL Interdisciplinary Postdoc Programme (EIPOD) under Marie Curie Actions COFUND (Y.L.), the 4D Nucleome / 4DN NIH Common Fund (U01 EB021223) (J.R.), the European Molecular Biology Laboratory (Y.L., E.B. J.V.R. and J.R.) and the startup grant from Southern University of Science and Technology (Y.L.).

## AUTHOR CONTRIBUTIONS

Y.L. E.B. and J.R. conceived the approach, developed the methods, wrote the software and analyzed the data. Y.Z. and J. B. contributed to the photometry-based data analysis. J.V.R. prepared the nuclear pore complexes sample. Y.L., and J. R. wrote the manuscript with the input from all other authors.

## Data availability statement

The datasets generated and analysed during the current study are available from the corresponding author upon reasonable request.

## Methods

### Sample preparation

Samples were imaged within a coverslip sandwich. This consisted of 2 coverslips one on top of another. The bottom coverslip had genome edited U20S Nup96-SNAP cells seeded on it. Beads were introduced to both the top and bottom coverslip to determine the thickness of the coverslip sandwich.

U2OS Nup96-SNAP cells were seeded on cleaned high-precision 24 mm round glass coverslips. The cells were seeded such that they were achieve 50-70% confluency on the day of fixation and staining. For fixation, the cells were first prefixed for 30 secs in fixation buffer (2.4% formaldehyde in PBS). They were then incubated for 3 mins in 0.4 % (v/v) Triton X-100 in PBS for permeabilization. After permeabilization, the cells were then incubated in fixation buffer for another 30 mins. The fixation buffer was then quenched using PBS containing 100 mM NH4Cl. To mitigate unspecific dye binding, the coverslip was incubated in Image-IT FX signal (ThermoFisher Scientific) enhancer for 30 mins. After the blocking step, the cells were stained with SNAP dye buffer (1 µM BG-AF647 (New England Biolabs, no. S9136S), 1 µM dithiothreitol, in 0.5 % (w/v) BSA in PBS) for 2 hours at room temperature. After staining, the coverslip was washed three times for 5 mins.

We then introduced beads to the coverslip. We performed an initial 200X dilution of TetraSpeck™ Microspheres, 0.1 µm (Invitrogen, T7279), into a thin-walled PCR tube (0.5 µL of beads into 100 µL of water). The diluted beads were then sonicated in a water bath sonicator at 4 °C for 5 mins. Then, we performed a serial dilution of the sonicated beads to get a final dilution of 20,000X. This new dilution was then subjected to sonication for 5 mins in a water bath sonicator at 4 °C. The beads were added to the coverslip together with 100 µL of 0.2 M MgCl2. To introduce beads to the top coverslip of the coverslip sandwich, we used 2 µL of sonicated 20,000X beads, and added them to 98 µL of water and onto the coverslip. To introduce beads to the bottom coverslip of the coverslip sandwich we used 10 µL of sonicated 20,000X beads and added them with 90 µL of water onto the coverslip. The bead solutions were incubated for 2 mins, after which both coverslips were rinsed twice with 3 mL of PBS.

2,2’-thiodiethanol was supplemented into the imaging buffer to achieve a refractive index, n = 1.40. The final composition of the imaging buffer consisted of 28.5 % (v/v) 2,2’-thiodiethanol (Sigma Aldrich, 166782-100G), 50 mM Tris/HCl pH 8, 10 mM NaCl, 10 % (w/v) D-glucose, 500 µg ml^−1^ glucose oxidase, 40 µg ml^−1^ glucose catalase and 35 mM MEA in water. The refractive index of the imaging buffer was determined using a refractometer.

After the final PBS rinse, the coverslip containing the cells was held vertically to allow residual PBS to drip dry. A Kimwipe tissue was used to wick away the final drop. The coverslip was then placed on a Kimwipe tissue with the cells facing up. This was done to ensure that the bottom surface of the coverslip is dried. 100 µL of refractive index matched imaging buffer was added to the bottom coverslip. The drying steps described earlier was also performed on the coverslip that was to be the top on the coverslip sandwich. Once dried, the top coverslip was gently lowed on to the bottom coverslip allowing the imaging buffer droplet to come in contact with the top coverslip. This was done to prevent air bubbles within the coverslip sandwich. The coverslips were then gently pushed against each other to remove excess imaging buffer. The excess imaging buffer was wicked away using a kimwipe tissue. Once most of the imaging buffer was removed. The coverslip sandwich was transferred onto the stage mount and sealed using 2 component silicon glue.

### IAB based 4Pi-PSF model

We represent the ideal 4Pi-PSF of a single emitter by coherently adding counter-propagating fields *E*_1_*(x, y, z)* and *E*_2_*(x, y, z)* from the upper and lower objectives, respectively. Assuming that the interference phase between the two fields is *φ*, the overall intensity of the coherent 4Pi-PSF can be written as

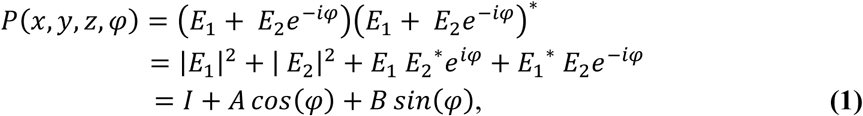

using the following substitutions:

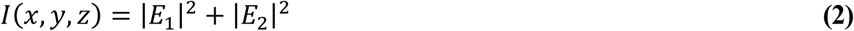

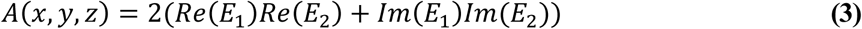

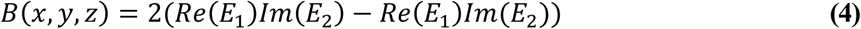

Here, *I(x, y, z), A(x, y, z)*and *B(x, y, z)*are slowly varying functions of *x, y, z*. They are real and same for all four quadrants. From Equation (1), we see that the three 3D matrices *I, A* and *B* are sufficient to fully describe the four-dimensional 4Pi-PSF, *P(x, y, z, φ).*

We call this model IAB-based 4Pi-PSF model. A fit with this model (see below) results in the coordinates *x, y, z* and the interference phase *φ*· *φ* depends on the OPD of the two arms and thus on the precise *z*-position of the fluorophore, but also has *z*-independent contributions *φ*_0_ (refractive index change, interference cavity drift induced OPD change, phase difference between quadrants, *etc*.): *φ* = 2*kz*_*φ*_ *+ φ*_0_.Usually, *φ*_0_ is slowly varying in time and similar for neighboring localizations and it causes an apparent sample *z* drift, which can be corrected by post processing. The *z*-position calculated from the interference phase can be written as: *z*_*φ*_ = *φ/*2*k.* Here, the wavenumber *K* = 2*nπ/λ.* and *n* is the refractive index of the immersion medium. It is worth to note that *z*_*φ*_ calculated from the interference phase is much more precise than the *z* parameter within the I, A, B matrices, which corresponds to the distance of the point emitter to the focal plane of the objective. Therefore, we use *z*_*φ*_ for the final reconstruction of the super-resolution image and use *z* for the phase unwrapping between different interference periods.

In order to determine the four-dimensional 4Pi-PSF experimentally, we need to determine the three 3D matrices *I, A* and *B*. They can be calculated from three experimental 4Pi-PSFs with known interference phase to solve for the three unknown parameters. Since matrix *I* is the sum of the intensity of the upper and lower single objective PSF, it can be easily obtained by different ways: direct sum of the 3D single objective PSFs from upper and lower objectives, sum of 2 interference 4Pi-PSF with the phase shift of *π* or sum of 4Pi-PSFs from all phase channels. Therefore, we actually only need two 4Pi-PSFs with known interference phase (phase shift other than *π*) to calculate the remaining two matrices. By solving two linear equations, we can obtain *A* and *B* :

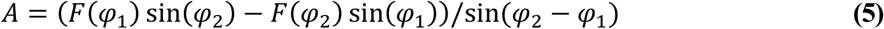

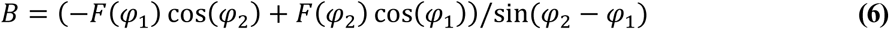

Here, *F(φ*_1_) = *P(φ* = *φ*_1_) − *I, F(φ*_2_) = *P(φ* = *φ*_2_) − *I* · *φ*_1_ and *φ*_2_ are the two interference phases of the two 4Pi-PSFs. Since different interference phase channels are usually acquired simultaneously in 4Pi-SMLM, it is easy to obtain two 4Pi-PSFs with different phases just by scanning the 4Pi-PSF in 3D once.

During the derivation of our new IAB-based 4Pi-PSF model, we did not use approximations, thus it can accurately describe even imperfect PSFs in presence of aberrations. We validated this by simulations using a full vectorial model to calculate the single-objective PSFs^19^, with different aberrations for upper and lower objectives, misalignment (lateral shift between PSFs) and astigmatism (**Supplementary Fig. S2**), often introduced to distinguish *z*-positions at different interference periods (‘phase unwrapping’)^8,20^. Furthermore, in order to be close to a realistic 4Pi-PSF, we added up the counter-propagating electrical fields from the upper and lower objectives partially coherently and incoherently^16^:

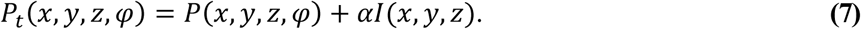

Here, *α* ≥ 0 is the ratio between the incoherent part and coherent part of the electrical field (*α* = 0.2 was used for simulation). The matrix I is the same as in **Equation (2)**. We then used the theoretical 4Pi-PSF model *P*_*t*_ to generate two 4Pi-PSFs with known interference phase (*φ*_1_ = 0, *φ*_2_ = *π/*2). Using **Equations (5)** and **(6)**, we could generate the A and B matrices. With all three parameters computed, we can generate 4Pi-PSFs with arbitrary phases using Equation (1). Here, we chose a phase of *φ* = 2*π/*3, which is different from the phases used to calculate I, A and B (*φ*_1_ = 0, *φ*_2_ = *π/*2) and compared it with the theoretical 4Pi-PSF with the same phase (**Supplementary Fig. S2**). As expected, these two PSFs are identical within machine precision.

### Fitting of multichannel 4Pi data

In 4Pi-SMLM experiments, three or four phases with relative phase shifts of *ϕ_i_* are detected simultaneously. To extract the maximum information in a robust way from these images of blinking fluorophores, we developed a global MLE algorithm to simultaneously fit the image of a single fluorophore in all channels. The imaging model of multi-channel 4Pi-PSF is given as:

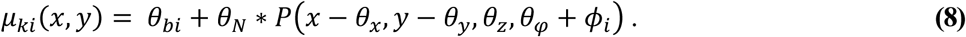

Here, *μ*_*ki*_ is the expected intensity value of pixel *k* at position *(x, y)* in channel *i* and is assumed to follow Poisson statistics due to the random nature of photon detection. For each channel, *θ*_*bi*_ are the background photons per pixel and assumed to be constant over the extent of the PSF in a given channel. *θ*_*N*_ is the number of the photons emitted from the single molecule. *θ*_*X*_, *θ*_*y*_ and *θ*_*z*_ are the *x, y* and *z* positions of the emitter. *θ*_*φ*_ is the interference phase. *P* is described as in Equation (1). The objective function for MLE across different channels is given by:

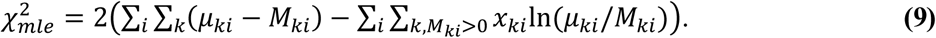

Here, *M*_*ki*_is the measured number of photons in the k^th^ pixel in the i^th^ channel. Similar to a previous implementation^21^, we used a modified Levenberg-Marquardt algorithm for the nonlinear optimization process. In order to calculate the partial derivatives along each parameter to construct the Hessian and Jacobian matrix during the iterative process, we used cubic spline interpolation to interpolate the 3D matrices *I, A* and *B*^15,22^. Since the interference phase is decoupled from the *θ*_*z*_ in the IAB-based 4Pi-PSF model, it is relatively simple to calculate the partial derivatives along each parameter (**Supplementary Note 1**).

During the global fitting procedure, the phase shifts *ϕ* _*i*_ between different channels can be precisely determined and do not change between experiments as they directly follow the design of the interference cavity. *θ*_*z*_ is also considered to be identical for all channels as the calibration at different *z*-positions is performed simultaneously for all channels.

