## Supplemental Material for "Accurate 4Pi single-molecule localization using an experimental PSF model"

**Supplementary Note 1. Levenberg-Marquart iterative schemes for MLE of IAB-based 4Pi-PSF model**

A modified Levenberg-Marquardt algorithm is used to minimize $\chi^2_{mle}$ (Equation (9) in main text). The iterative process is similar as in (ref. 1) where Hessian and Jacobian matrix was used to calculate the update of parameters in each iteration. Here, the second-derivatives term is neglected and only the first derivatives are used (ref. 1). In order to construct the Hessian and Jacobian matrix, the following partial derivatives are used:

$$\frac{\partial\mu_{ki}}{\partial\theta_x} = \theta_N\left(\frac{\partial \mathrm{I}(x-\theta_x,y-\theta_y,\theta_z)}{\partial\theta_x} + \frac{\partial \mathrm{A}(x-\theta_x,y-\theta_y,\theta_z)}{\partial\theta_x}\cos(\theta_\varphi+\phi_i) + \frac{\partial \mathrm{B}(x-\theta_x,y-\theta_y,\theta_z)}{\partial\theta_x}\sin(\theta_\varphi+\phi_i)\right), \qquad \textbf{(S1)}$$

$$\frac{\partial\mu_{ki}}{\partial\theta_y} = \theta_N\left(\frac{\partial \mathrm{I}(x-\theta_x,y-\theta_y,\theta_z)}{\partial\theta_y} + \frac{\partial \mathrm{A}(x-\theta_x,y-\theta_y,\theta_z)}{\partial\theta_y}\cos(\theta_\varphi+\phi_i) + \frac{\partial \mathrm{B}(x-\theta_x,y-\theta_y,\theta_z)}{\partial\theta_y}\sin(\theta_\varphi+\phi_i)\right), \qquad \textbf{(S2)}$$

$$\frac{\partial\mu_{ki}}{\partial\theta_z} = \theta_N\left(\frac{\partial \mathrm{I}(x-\theta_x,y-\theta_y,\theta_z)}{\partial\theta_z} + \frac{\partial \mathrm{A}(x-\theta_x,y-\theta_y,\theta_z)}{\partial\theta_z}\cos(\theta_\varphi+\phi_i) + \frac{\partial \mathrm{B}(x-\theta_x,y-\theta_y,\theta_z)}{\partial\theta_z}\sin(\theta_\varphi+\phi_i)\right), \qquad \textbf{(S3)}$$

$$\frac{\partial\mu_{ki}}{\partial\theta_\varphi} = \theta_N\left(-\mathrm{A}(x-\theta_x,y-\theta_y,\theta_z)\sin(\theta_\varphi+\phi_i) + \mathrm{B}(x-\theta_x,y-\theta_y,\theta_z)\cos(\theta_\varphi+\phi_i)\right), \qquad \textbf{(S4)}$$

$$\frac{\partial\mu_{ki}}{\partial\theta_N} = \mathrm{I}(x-\theta_x,y-\theta_y,\theta_z) + \mathrm{A}(x-\theta_x,y-\theta_y,\theta_z)\cos(\theta_\varphi+\phi_i) + \mathrm{B}(x-\theta_x,y-\theta_y,\theta_z)\sin(\theta_\varphi+\phi_i), \qquad \textbf{(S5)}$$

$$\frac{\partial\mu_{ki}}{\partial\theta_{bi}} = 1 \qquad \textbf{(S6)}$$

Here, the cubic splines are used to interpolate the 3D matrices I, A and B to calculate the partial derivative along x, y and z, separately [1, 2]. The iterative process is considered to be converged when the ratio of the relative change of $\chi^2_{\mathrm{mle}}$ is less than $10^{-6}$ compared to the last iteration.

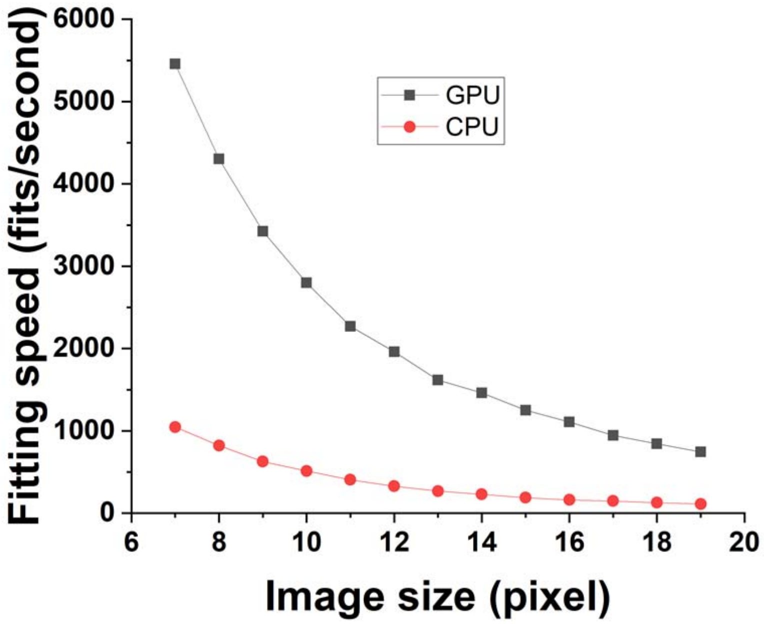


**Figure S1. Computational speed of the fitting routines in GPU and CPU.** A consumer graphic card NVIDIA GeForce GTX 1070 was used for GPU calculation. Intel Core i7-5930 was used for CPU calculation. The size of fitting region was chosen from 7×7 to 19×19 pixels.

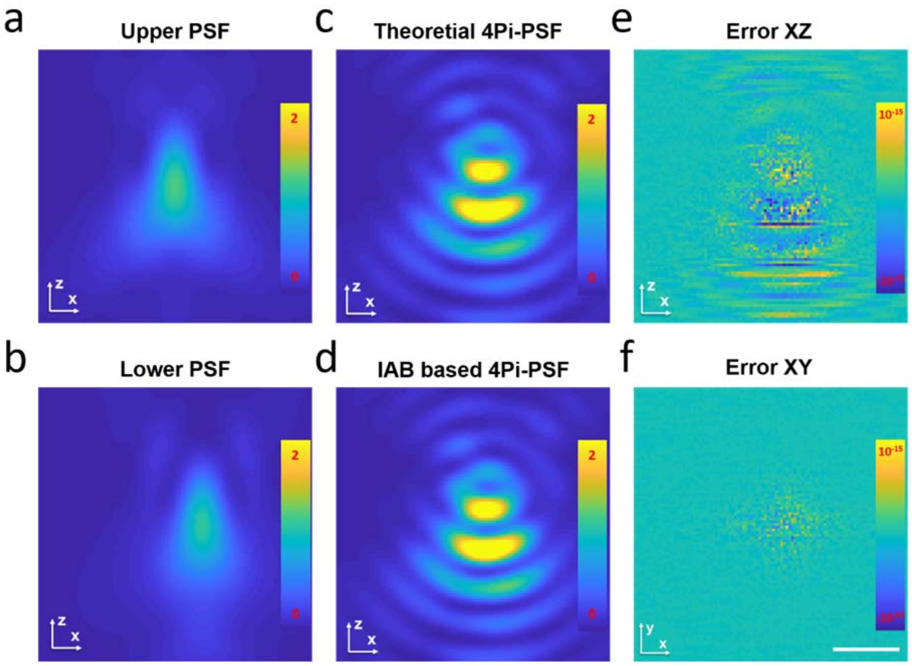


**Figure S2. Comparison of IAB based 4Pi-PSF model and theoretical 4Pi-PSF model.** (a) Upper objective PSF with astigmatism (100 mλ) and spherical (20 mλ) aberrations. (b) Lower objective PSF with astigmatism aberration (-100 mλ). The PSF was shifted in x by 200 nm. (c) Theoretical 4Pi-PSF contains both coherent and incoherent part. The ratio between the ratio between the incoherent part and coherent part $\alpha$ is 0.2. (d) IAB model based 4Pi-PSF calculated using Equation (1). (e) x-z cross section of the error between theoretical 4Pi-PSF and IAB model based 4Pi-PSF. (f)The error between theoretical 4Pi-PSF and IAB model based 4Pi-PSF in xy. Scale bar 500 nm.
